# Temporal changes in genetic admixture are linked to heterozygosity and health diagnoses in humans

**DOI:** 10.1101/697581

**Authors:** Brian S. Mautz, Jacklyn N. Hellwege, Chun Li, Yaomin Xu, Siwei Zhang, Joshua C. Denny, Dan M. Roden, Tracy L. McGregor, Digna R. Velez Edwards, Todd L. Edwards

## Abstract

Reproduction between individuals from different ancestral populations creates genetically admixed offspring. Admixture can have positive and negative impacts on individual health, feeding back to population health. Historical and forced migrations, and recent mobility, have brought formerly disparate populations of humans together. Here we sought to better understand how temporal changes in genetic admixture influence levels of heterozygosity and health outcomes. We evaluated variation in ancestry over 100 birth years in 35,842 individuals from a genetic database linked to health records in a population in the Southeastern United States. Analysis of 2,678 ancestrally informative markers revealed increased admixture and heterozygosity for all clinically-defined race groups since 1990. Most groups also exhibited increasing long-range linkage disequilibrium over time. A phenome-wide association study of clinical outcomes detected protective associations with female reproductive disorders and increased risk for diseases with links to autoimmunity dysfunction. These mixed effects have important ramifications for human health.

## Introduction

Understanding temporal changes in heterozygosity has important implications for individual- and population-level health in humans that remain unexplored. The heterozygosity-fitness correlation (HFC) is a measure used to quantify the extent to which proxies of fitness are positively correlated with the level of heterozygosity across sets of genetic loci or the whole genome (David, 1998; Head et al., 2017; Szulkin et al., 2010). Evidence linking fitness components to levels of heterozygosity has been found in animals (Keller and Waller, 2002), but results are equivocal and controversial (Chapman et al., 2009; Szulkin et al., 2010). HFCs arise through some level of inbreeding depression or can be generated through immigration and population admixture (reviewed in Szulkin et al., 2010). Both positive and negative effects on individual and population health can occur.

Three factors make human population genetic data ideal to study the connection between heterozygosity and health. First, the human genome is sequenced and genetic databases with large numbers of genetic loci and individuals are readily available. Second, there are substantial numbers of subjects with genetic data linked to electronic health records (EHR) (Roden et al., 2008; Sudlow et al., 2015). Third, the diseases and their etiologies used in health records are known in much more detail and well categorized than most other species, facilitating the estimation the relationship between heterozygosity and changes in (disease) phenotypes used as fitness proxies.

Additionally, we have detailed information on the movements of many human populations. Geographically disparate human populations have come into contact through migration, as well as more recent mobility and urbanization, increasing the opportunity for genetic admixture and HFCs to occur. Genetic admixture has previously been used to identify migration patterns across several humans populations (Baharian et al., 2016; Bryc et al., 2010, 2015; Han et al., 2017; Moreno-Estrada et al., 2014; Tishkoff et al., 2009; Wang et al., 2013) and the genetic basis of diseases (Franceschini et al., 2013; Kato et al., 2015; Reich and Patterson, 2005). Two studies have shown temporal increases in heterozygosity due to urbanization, one in a Croatian population (Rudan et al., 2008) and one in a US population of European ancestry (Nalls et al., 2009). However, these studies of admixture have not connected migratory or urbanization patterns to health outcomes.

Finally, accruing evidence in humans suggests HFCs occur. For example, several studies have linked levels of heterozygosity in non-admixed populations to specific disease states (Bittles and Black, 2010; Campbell et al., 2007, 2009; Pemberton and Szpiech, 2018). Additionally, a meta-analysis of populations of European American and African American ancestry found a positive association between levels of heterozygosity and mortality in humans (Bihlmeyer et al., 2014). However, these studies are limited in scope, investigating single outcomes and ignoring the impact of temporal trends in admixture and heterozygosity on epidemiological outcomes.

Here, we link temporal changes in genetic admixture to patterns of heterozygosity and epidemiological outcomes across a broad array of human diseases. We used BioVU, a biorepository of genetic data linked to EHR (Roden et al., 2008) to investigate temporal changes in genetic admixture and heterozygosity in a population in the Southeastern United States. Though race has little biological meaning, we used race as instrumental groupings (Maglo et al., 2016) to reflect corresponding socio-cultural dynamics in this geographic area and investigate differential changes to admixture and heterozygosity within these groupings. In addition, we used a phenome-wide association study (PheWAS, Denny et al., 2010), to connect genetic data with the clinical phenome representing an entire collection of disease outcomes associated with EHRs in BioVU.

## Results

We identified 2,678 ancestry-informative markers (AIMs) from genetic data in BioVU. These AIMs were from ExomeChip data in a cohort of 28,723 Whites, 4,129 Blacks, 550 Hispanic/Latinos, and 270 Asians, based on the EHR third-party race designation (see methods). Sample population demographics by EHR race and birth decade are in presented in Supplementary Table S1-2 and Figure S1. After combining our data with the 1000 Genomes as a reference group (Durbin et al., 2010), we calculated principal components to identify patterns of ancestry in each individual. We used the ancestral classifications to test for temporal trends in mean and variance in admixture proportion.

Analysis of temporal trends in genetic admixture showed an increase in ancestral diversity with time (Figure 1; see Table S2). The admixture proportion consistently increased with younger ages in Asian and Hispanic/Latino groups. In Whites, the level of admixture remained stable until approximately 1990 when it began to increase. Blacks had a decrease in admixture proportion in those born the early- to mid-20th century, but an increase after the 1980’s. In addition to the distinct clusters in the principal component plots representing individuals who are identified as either White and Black and who contain both European and African ancestries, the overall proportions of ancestry across all recorded race groups exhibit increasing admixture in younger individuals (Figure S3-S7). Additionally, a significant increase in the variance of admixture proportions over time was observed (variance coefficient = 0.0193 ± 0.0009 [SE], p-value < 1.44×10^−100^), indicating that there is a linear increase in variance of the admixture proportion of 0.0193 with every birth year. This is consistent with recent and ongoing admixture (Verdu and Rosenberg, 2011).

**Figure 1.**
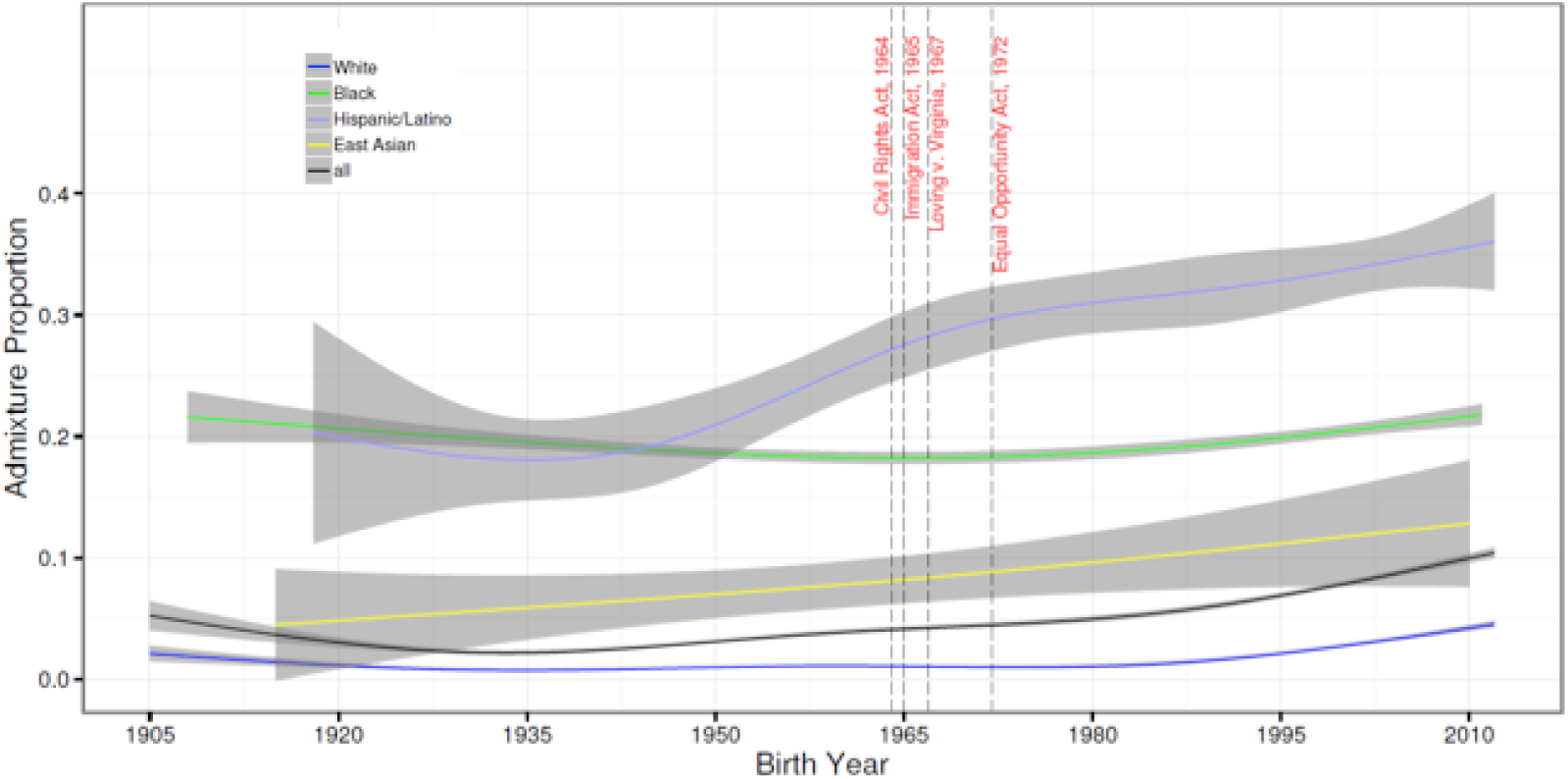
Admixture proportion plotted against birth year. Admixture proportion is defined as (1-maximum ancestry proportion). The smoothing curves are obtained using the generalized additive model method with a cubic spline basis implemented in R package mgcv and plotted using R package ggplot2. Cohorts are grouped by race and 95% confidence intervals are included.

We detect similar patterns in three additional sources. First, we calculated heterozygosity across AIMs and tested for temporal trends to compare with patterns of genetic admixture. There was strong support for increasing heterozygosity in younger individuals (Figure 2; Table S3 and Figure S8). The timing of inflection for increased standardized heterozygosity varied between the race groups, but the data indicated that ancestry diversity has accelerated rapidly in Asian, Black, and White cohorts since about 1980, while Hispanics/Latinos have exhibited a relatively steady rate of increasing diversity since the 1940s. This reflects the increasing number of children born to couples of predominantly different ancestral backgrounds over the past few decades.

**Figure 2.**
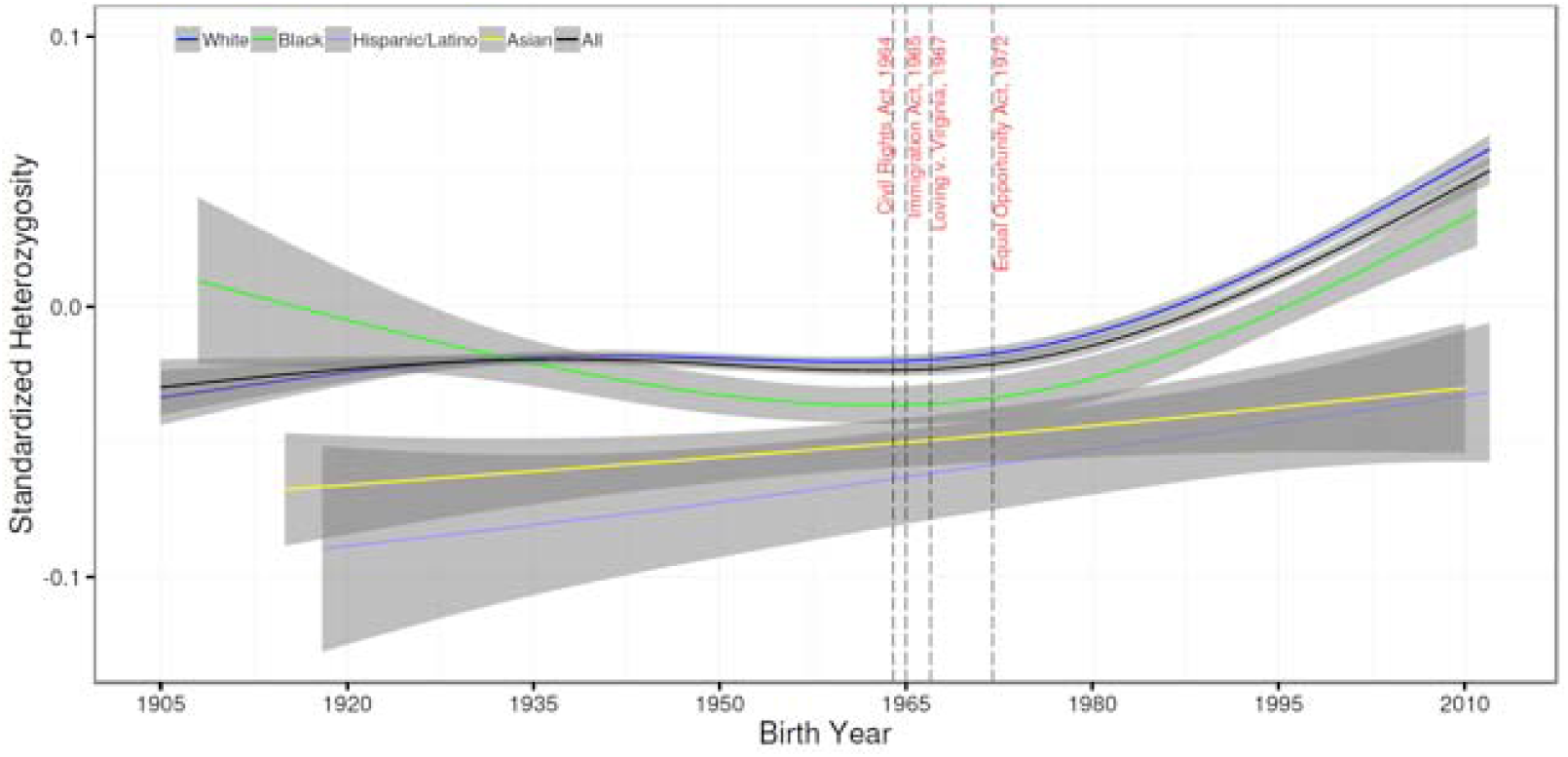
Standardized heterozygosity plotted against birth year. The y-axis depicts the standardized heterozygosity of individuals with birth year on the x-axis. Cohorts are grouped by stated race and 95% confidence intervals are indicated.

Second, we verified patterns by estimating pairwise long-range linkage disequilibrium in our dataset. Using common single nucleotide polymorphisms (SNPs) (minor allele frequency [MAF] > 5%) from the genotyping array, we calculated pairwise D’ using Haploview (Barrett et al., 2005) and plotted D’ against physical distance for all pairs of SNPs between 9–10 megabases of each other (Figure 3; Table S4). Results show a small drop in long range disequilibrium (LRLD) in individuals born before the 1950s, followed by significant steady increases in the 1950s through the 1980s and subtle fluctuation at higher levels in the 1990s to 2010s.

**Figure 3.**
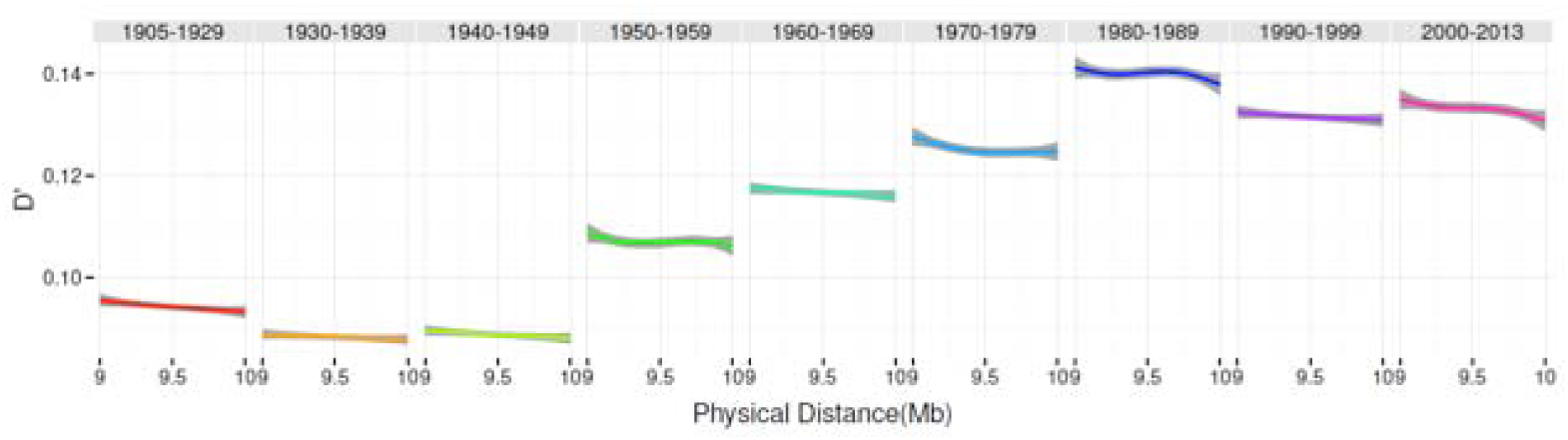
Pairwise D’ for SNPs between 9-10 Mb for all common SNPs on the exome array in all study individuals stratified by intervals of birth decade. Pairwise D’ for all common SNPs between 9 - 10 megabases apart for individuals are presented on the y-axis and physical distance in Mb is presented on the x-axis. The plots are paneled by decades of birth to depict relative changes over time.

Third, we estimated changes in the number of ethnicities in self-reported ancestry in the 2013 American Communities Survey. Respondents were instructed to select Hispanic/Latino/Spanish status and all applicable races for each individual in the household (Ruggles, 2014). We analyzed the average number of race categories selected for each individual by birth year and stratified these results within race categories for Tennessee (Figure S9), the East South-Central Region which includes Tennessee, Alabama, Kentucky, and Mississippi (Figure S10), and the entire United States (Figure S11). The results from all regions are concordant, indicating that younger individuals are more likely to indicate multiple races. The inflection point appears to have been earlier in the Asian race category but is demonstrated in all race groups by the mid-1960s in the entire US sample.

### Changes in health diagnoses with admixture

We evaluated the association between individual heterozygosity and disease diagnoses using a phenome-wide association study approach (PheWAS, Denny et al., 2010). Increasing genetic admixture resulted in fewer diagnoses of female reproductive traits across all data (Table 1). These results remain statistically significant when considering adults only (Table S5) and after correction for multiple tests. Phenotype codes for “disorders of menstruation and other abnormal bleeding from female genital tract” and “irregular menstrual cycle/bleeding” were significantly associated with protection by increasing heterozygosity (p-value = 7.21×10^−6^ and 4.37×10^−5^, respectively; Table 1, Figure S12). Other protective findings were also gynecological in nature, including cervical cancer/dysplasia and abnormal Papanicolaou smear results. Significant phenotypes in adults were predominantly detected for females. Outside genitourinary findings, other nominally significant associations (Bonferroni significant < *p* ≤ 0.05) show increased risk with genetic admixture and include atopic dermatitis, AV Block, obstructive asthma, Sicca syndrome.

**Table 1.** Results from the phenome-wide association study of heterozygosity and clinical outcomes for full sample.

| PheCode | Phenotype | P-value | OR (95% Confidence Interval) |
| --- | --- | --- | --- |
| 626 | Disorders of menstruation and other abnormal bleeding from female genital tract | $7.21 \times 10^{-6}$ | 0.37 (0.24 – 0.57) |
| 626.1 | Irregular menstrual cycle/bleeding | $4.37 \times 10^{-5}$ | 0.37 (0.23 – 0.60) |
| 939 | Atopic/contact dermatitis due to other or unspecified | $2.86 \times 10^{-4}$ | 1.82 (1.32 – 2.52) |
| 180 | Cervical cancer and dysplasia | $4.02 \times 10^{-4}$ | 0.19 (0.08 – 0.48) |
| 792.1 | Papanicolaou smear of cervix or vagina with atypical squamous cells | $5.36 \times 10^{-4}$ | 0.21 (0.09 – 0.51) |
| 180.3 | Cervical intraepithelial neoplasia [CIN] [Cervical dysplasia] | $6.24 \times 10^{-4}$ | 0.15 (0.05 – 0.45) |
| 426.2 | Atrioventricular [AV] block | $7.62 \times 10^{-4}$ | 2.97 (1.58 – 5.61) |
| 495.11 | Chronic obstructive asthma with exacerbation | $8.13 \times 10^{-4}$ | 4.63 (1.89 – 11.34) |
| 785 | Abdominal pain | $1.04 \times 10^{-3}$ | 0.70 (0.57 – 0.87) |
| 792 | Abnormal Papanicolaou smear of cervix and cervical Human Papilloma Virus (HPV) | $1.04 \times 10^{-3}$ | 0.33 (0.17 – 0.64) |
| 709.2 | Sicca syndrome | $1.66 \times 10^{-3}$ | 4.38 (1.75 – 10.99) |
| 318 | Tobacco use disorder | $1.75 \times 10^{-3}$ | 0.48 (0.30 – 0.76) |
| 626.12 | Excessive or frequent menstruation | $1.80 \times 10^{-3}$ | 0.28 (0.12 – 0.62) |
| 628 | Ovarian cyst | $2.83 \times 10^{-3}$ | 0.23 (0.09 – 0.60) |
| 426.24 | Atrioventricular block, complete | $3.57 \times 10^{-3}$ | 3.30 (1.48 – 7.38) |

## Discussion

We found increasing genetic admixture and heterozygosity through time in a population of the Southeastern United States. Comparing these changes to socio-cultural shifts in this geographic area puts these results in context. In the Southern United States from the Civil War until the mid-1960’s, laws and policies enforced segregation of populations of European and African ancestry. Consistent with these socio-cultural boundaries, there is little change in admixture through the 1960’s. Additionally, despite legal rulings and perceived socio-cultural transformation, there remained a very slow increase in admixture and heterozygosity for an additional 20-30 years. For individuals with an EHR designation of White, the mean proportion of European ancestry was greater than 98% until the 1990’s but decreased to 92% in the youngest cohort as the proportion of African ancestry increased to 6% (Figure S4). In individuals designated as Black in the EHR, the mean African proportion was approximately 80% in those born before the 1990’s but decreased to 77% in those born more recently (Figure S5). The largest change of any group was in those identified as Hispanic/Latino. The European ancestry proportion in these individuals decreased by 15%, from approximately 75% in those born before the 1950’s to under 60% after the 1980’s. This decrease in European ancestry was associated with an increase in the East Asian ancestry proportion, likely a proxy for Native American ancestry, while the African proportion remained under 10% (Figure S6). The continental ancestry proportions in EHR-designated Asians were more difficult to discern because of the small cohort size, although we did observe a slight decrease in the East Asian proportion over time (Figure S7). These trends are consistent with previous reported trends in decreases in homozygosity in other populations (Nalls et al., 2009; Rudan et al., 2008).

Long-range linkage disequilibrium occurs when subdivided populations mix (Koch et al., 2013; Nei and Li, 1973). Consistent with this and with patterns we detected in admixture and heterozygosity, we find that LRLD changes through time (Figure 3). There was small drop in LRLD for older individuals born before 1950. This was followed by a consistent increase in LRLD in individuals born up to 1980. Subtle fluctuations at higher LRDL levels occurs in those individuals born between 1990 and 2010. It should be noted, however, that we cannot necessarily rule out other possible sources of LRLD (e.g. drift, epistatic selection, hitchhiking as in Koch et al., 2013), though they seem unlikely given the recent nature of admixture and formation of LRLD.

Our results are qualitatively echoed in the 2013 American Communities Survey. Younger individuals, across multiple geographic areas (Tennessee, East South-Central Region, United States) are more likely to indicate multiple races when asked about their ancestral origins (Figure S9-11). Across race group the inflection point occurs by the mid-1960’s of the entire US sample, though appears earlier for the Asian race category.

### Changes in health diagnoses with admixture

We show that changes in population genetic parameters have important consequences for population health (Table 1; Table S5 and Figure S12), which have not been previously documented. There were several statistically significant associations of genetic admixture with adult female genitourinary diagnosis codes (e.g. irregular menstrual cycle/bleeding; cervical cancer/dysplasia). These findings linking admixture to protection from menstruation and gynecological abnormalities suggests that ancestral diversity is related to risk of disorders that could affect reproduction. This is consistent with long-standing models that propose ancestral diversity and heterozygosity is related to fertility in plants and animals, as well as other beneficial characteristics in offspring (Dickerson, 1973; Keller and Waller, 2002; Lamkey and Edwards, 1999; Shull, 1948; Szulkin et al., 2010).

The changes in reproductive diagnoses were detected predominantly for females, suggesting a largely sex-specific response to changes in admixture and heterozygosity. In some animals, there are sex-specific responses in reproduction to inbreeding (Keller and Waller, 2002), providing some circumstantial support for our findings. However, the sex-specific response we detected could also be a results of gender differences in other factors. For example, men are less likely to seek medical attention and report medical issues (Galdas et al., 2005; Mansfield et al., 2003). Alternatively, male reproductive traits, such as sperm quality, might not be routinely checked and reported as with female reproductive parameters. More research is needed to disentangle biological and sociological factors driving the sex-specific responses to determine the source of these trends.

Other interesting patterns emerge when considering all data and nominally significant PheWAS results (i.e. p < 0.05). Several of these disease phenotypes, show positive associations with increasing genetic admixture. Importantly, each of these diseases have at least suggestive links to autoimmunity issues (*atopic dermatitis*: Mittermann et al., 2004, AV block: Buyon et al., 1998; Villuendas et al., 2014, asthma: Barnes, 2008; Tedeschi and Asero, 2008, *Sicca/Sjögren syndrome*: Brito-Zerón et al., 2016). The nominally significant diseases detected here also have reported connections with each other in the literature. For example, atopic dermatitis and asthma have a similar mechanistic basis via allergen exposure through the epidermis (Spergel, 2010). Similarly, autoimmune congenital heart block is linked to Ro and La autoantigens, Sjögren-syndrome-related antigen A and B respectively, which cross the placenta from the mother to the fetus to cause cardiac abnormalities and cutaneous rashes (Brito-Zerón et al., 2015). These patterns suggest increasing heterozygosity are linked to increases in activity in the immune system. Our results show continued increases in genetic admixture with time. If heterozygosity is positively linked to autoimmunity-related disease, then we would predict continued increases in prevalence of these types of disease with time. Future research should address these intriguing immunity-disease relationships to determine the validity and consistency of these patterns within and across human populations.

These results have important ramifications in several areas. First, this work has implications for genetic association study designs, disease prevalence and disparities, and classification of individuals for their medical care. Frequently genetic analyses are stratified by ancestry or race, and our data show that simple categorical race classifiers, whether by self-report or derived from other data, are becoming less accurate over time. When conducting genetic studies in children, researchers may need to collect the race(s) of the parents or even grandparents to allow for appropriate classification, or consider AIMs to estimate ancestry proportions (Kodaman et al., 2013; Ruiz-Narvá Ez et al., 2011). This increase in ancestral diversity of the younger population may also enable admixture mapping studies in populations not traditionally considered for these types of studies.

Second, understanding race-associated factors in patients with complex ancestries may be increasingly important for effectively delivering precision medical care, which is a major National Institutes of Health (NIH) goal and one of the key promises of the genetics research enterprise in the United States. Genetic information has been used to investigate the roots of human evolution and subsequent diversity. These same approaches can be applied to analyze ongoing trends in regional and national demographics which have significant implications for scientific, societal, and governmental policies.

Finally, these results have important implications for HFCs and their use in biological research (Keller and Waller, 2002; Chapman et al., 2009; Szulkin et al. 2010). We detected positive changes in reproductive traits in females, providing evidence that genetic admixture and heterozygosity are linked to potential fitness-related components. Though there were protective effects for reproductive diagnoses, our results provide additional details showing increased risk associated with other disease states that might be linked with the immune system as genetic admixture increases and that HFCs might be sex-specific. The effect of changes to levels on health and fitness related parameters are complex, so HFCs should be interpreted with caution and more detail is needed for proper interpretation.

## Materials and Methods

Individuals were selected from the BioVU DNA Repository which links clinical data from de-identified electronic medical records to DNA specimens obtained from patients at Vanderbilt University Medical Center (VUMC) (Roden et al., 2008). Each individual is assigned a single race in the Electronic Heath Record (EHR) of White, Black, Asian, Pacific Islander, American Indian/Alaska Native, or declined/unknown, and an ethnicity of Hispanic/Latino, Not Hispanic/Latino, or declined/unknown. BioVU also contains third-party-designated race, which is a good predictor of self-reported ethnicity/ancestry in this database (Dumitrescu et al., 2010). This study of de-identified data was determined to be non-human subject research by the institutional review board (IRB) of Vanderbilt University, Nashville, TN.

### DNA Extraction and Genotyping

All DNA samples were isolated from whole blood using the Autopure LS system (QIAGEN Inc., Valencia, CA). Genomic DNA was quantitated via an ND-8000 spectrophotometer and DNA quality was evaluated via gel electrophoresis. Individuals were genotyped using the Illumina Infinium HumanExome Array [12v1-1] (Illumina Inc., San Diego, CA). The data were processed for genotype calling using Illumina’s Genome Studio (Illumina Inc., San Diego, CA).

### Genotyping Quality Control

Data on 240,117 SNPs and 35,842 individuals (16,289 males, 19,552 females) were available prior to implementation of quality control (QC) measures. No individuals were excluded for low genotyping efficiency (<98%). 6,599 SNPs were excluded for low genotyping efficiency (<98%) and 71,667 SNPs were monomorphic. Twenty-six individuals (14 EHR males, 12 EHR females) were excluded for inconsistent genetic and database sex. After QC, 163,135 SNPs remained for analyses in 35,456 individuals. No SNPs were removed for deviations from Hardy-Weinberg equilibrium.

### Quantification and Statistical Analyses

Descriptive statistics of demographic and clinical characteristics were expressed as mean with standard deviation or median with interquartile range for continuous covariates and as frequencies or proportions for categorical data using SPSS statistical software (IBM Corporation, Armonk, NY).

A subset of 2,678 ancestry-informative markers (AIMs) were selected for subsequent analysis. We selected AIMs from the ExomeChip content, which were selected to have strong differences between African and European ancestry populations as well as Native American and European ancestry populations. Average differences in allele frequency, Δ_*p*_, between 1000 Genomes (Price et al., 2006) continental populations (European, Asian, Americas, African) for the AIMs are presented in Table S6. AIMs were used rather than using pruned SNP data due to the genotype data coming from an ExomeChip platform, which was designed with a panel of AIMs to enable evaluation of ancestry.

EIGENSTRAT v6.0.1 software was used to conduct principal components analysis (PCA) to estimate continuous axes of ancestry from AIMs running all populations together (Price et al., 2006).

STRUCTURE software v2.3.3 (Falush et al.; Pritchard et al., 2000) was used to quantify ancestry in combined study and 1000 Genomes Project (Price et al., 2006) individuals using the AIMs. We estimated proportions of ancestry assuming ancestral clusters (*K*) ranging from one to 16, where 16 is the number of sub-populations in the 1000 Genomes data plus two. We assumed unlinked SNPs and used 5,000 iterations of burn-in and 10,000 iterations for analysis without providing population information to the software. We observed that the –log-likelihood of the data given *K* did not vary significantly for *K*’s greater than three and observed that *K*’s greater than three primarily subdivided the European populations (data not shown). The three STRUCTURE clusters corresponded to African, European, and Asian ancestry based on comparisons to the 1000 Genomes reference data.

Each of the three derived proportions of continental ancestry from STRUCTURE were regressed onto birth year using generalized additive models with integrated smoothness estimation (GAM) (Wood, 2010) implemented in the R package mgcv in all study individuals, Non-Hispanic Whites, Non-Hispanic Blacks, Hispanics/Latinos, and Non-Hispanic Asians. We derived admixture proportion, *α*, for the *i*th individual using the formula *α*_*i*_ = 1 – maximum (%European, %African, %East Asian). We regressed *α*_*i*_ onto birth year using generalized additive models with integrated smoothness estimation for all study individuals, Non-Hispanic Whites, Non-Hispanic Blacks, Non-Hispanic Asians, and Hispanics or Latinos. Proportions of continental ancestry in the Non-Hispanic Whites, Non-Hispanic Blacks, Non-Hispanic Asians, and Hispanics or Latinos by birth decade are presented.

It has been previously shown by Verdu and Rosenberg (2011) that when parental populations stop contributing to admixture, that the variance of admixture proportions decreases rapidly, and when they continue to contribute to the admixture, that the variance of admixture proportions increases over time. To test null hypothesis that the observed levels of admixture in our data were not due to ongoing and increasing rates of admixture, we analyzed the association between the variance of *α* and birth year. We used the software package MVtest and modeled the admixture proportion variance as a log-linear function of birth year and five principal components of ancestry using estimating equations.

### Analysis of admixture proportion variance over time

We model admixture proportion mean and variance simultaneously as functions of birth year and covariates. Specifically, let *m*_*i*_ be the mean and 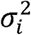 be the variance of the trait *Y*_*i*_ for the *i*-th individual. We model them as

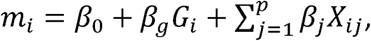

And

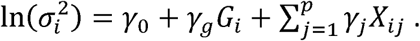

where *G*_*i*_ is the birth year or variable of interest for the *i*-th individual, and *X*_*i*1_, …, *X*_*ip*_ are *p* covariates. In this model the variance is monotonic with respect to birth year, an assumption that holds in most circumstances. The parameters are estimated simultaneously. This framework allows for testing of the null hypothesis of no effect on mean, variance, or both for any term in the model. These correspond to a mean test with null H_0_: *β*_*g*_ = 0, a variance test with H_0_: *γ*_*g*_ = 0, both having one degree of freedom (DF), and a 2-DF test with H_0_: *β*_*g*_ = 0, *γ*_*g*_ = 0.

### Model fitting with estimating equations

The parameters are estimated through the estimating equations approach, which does not require a full specification of the outcome distribution, but only a few constraints for the parameters of interest. These constraints are often written as equations, and the parameter estimates can be obtained by solving the equations. The asymptotic distribution for the parameter estimates can be derived (Stefanski and Boos, 2002). Specifically, suppose random variable *y*_*i*_ has mean 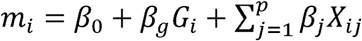, and log-variance ln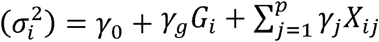. There are *k* = 2(*p* + 2) the parameters, which can be written as a vector ***θ*** = (***β, γ***),where ***β=*** (*β*_*0*_, *β*_*g*_, *β*_1_ *…,β*_*p*_) and ***γ=*** (*γ*_*0*_, *γ*_*g*_, *γ*_1_ *…,γ*_*p*_). Let *y*_*i*_ and ***x***_*i*_ = (1,*g*_*i*_,*x*_*i*1_, …, *x*_*ip*_)^*T*^ be the observed values for subject *i*. If we had assumed normality for the outcome, the log-likelihood for observation *i* would have been

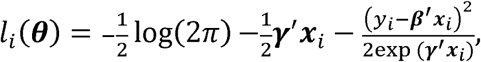

for which the partial derivatives with respect to the parameters ***θ*** is a *k*-vector,

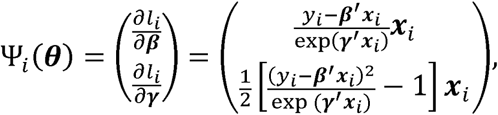

and the maximum likelihood estimates of the parameters could have been obtained by solving the *k* equations 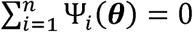. This m otivated us to use these *k* equations,

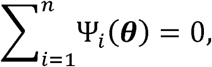

as the starting point for our estimating equations approach to obtain parameter estimates 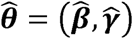. If normal ity holds, then 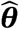 are the maximum likelihood estimates. Note that although the estimating equations were motivated by the Gaussian likelihood, one can always start from these equations to obtain 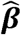 and 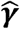, whether normality holds or not, and proceed with statistical inference using the M-estimation theory (Stefanski and Boos, 2002). This is a major advantage for using estimating equations. The partial derivative of Ψ_*i*_(***θ***) is a *k* × *k* matrix, denoted as *Ψ*_*i*_(***θ***). Using the M-estimation theory, we have

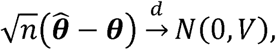

where the *k* × *k* covariance matrix *V* can be estimated as *A*^−1^ *B*(*A*^−1^)^*T*^ with

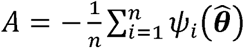 and 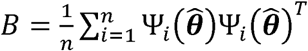.

If our interest is on the effect of *G*, the asymptotic result for the joint distribution for the parameter estimates 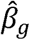 and 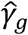 is

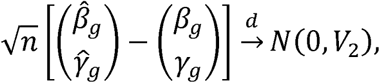

where *V*_2_ is the corresponding 2×2 submatrix of *V*, with diagonal values denoted as 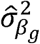 and 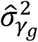, respectively. A mean test (H_0_: *β*_*g*_ = 0) can be performed by comparing 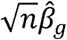 with 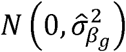, and similarly, a variance test (H_0_: *β*_*g*_ = 0) by comparing 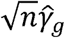 with 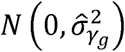. A 2-DF joint test (H_0_: *β*_*g*_ = 0, *γ*_*g*_ = 0) can be performed by comparing 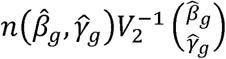 with a chi-squared distribution with two degrees of freedom. MVtest software for genetic analysis of SNP data or general analysis of variables is freely available at https://github.com/edwards-lab/MVtest.

### Heterozygosity Analysis

Standardized measures of heterozygosity among the AIMs were calculated in order to evaluate trends in heterozygosity over time relative to expectations. We first estimated the expected number of heterozygous genotypes in an individual in the k-th subpopulation as

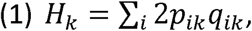

where the sum is over all SNPs in our analysis, and *p*_*ik*_ and *q*_*ik*_ = 1-*p*_*ik*_ are the allele frequencies for the *i*-th SNP. Hardy-Weinberg equilibrium was assumed. Then for every individual *j* in the *k*-th subpopulation, we standardized the observed number of heterozygous genotypes, *O*_*kj*_, by comparing it with the expected number *H*_*k*_:

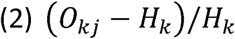

Standardized heterozygosity was regressed onto birth year using GAM in all study individuals, and the results were plotted for Non-Hispanic White, Non-Hispanic Black, Non-Hispanic Asian, and Hispanic or Latino.

Pairwise linkage disequilibrium (LD) D’ statistics were calculated for all pairs of common (MAF > 0.05) SNPs within 10 megabases (Mb) using Haploview software (Wood, 2011). D’ statistics were regressed onto physical distance between SNPs using GAM for distances in the interval from 9-10 Mb for each birth decade.

We downloaded a 1% representative sample of individual-level responses to the American Community Survey (ACS) from the Integrated Public Use Microdata Series (IPUMS) (Ruggles, 2014). Race and ethnicity was self-reported in the ACS IPUMS. We regressed the number of major race groups claimed by individuals onto their reported birth year using GAM and frequency weights provided by IPUMS for TN, the South East Central census region, and the entire US. For individual groups, such as “White” and “Black or African American”, we plotted all individuals who responded affirmatively to those items; thereby, the samples for the individual race group plots are not independent and overlap at observations where participants claim two or more race groups.

### Phenotype Classification

Each individual was classified according to 1,645 phenotypes based on the International Classification of Disease, Ninth Revision, Clinical Modification (ICD9) Codes (Denny et al., 2013). Our classification strategy includes all ICD9 features except for procedures. Additionally, the system is hierarchical such that disease subtypes are also classified, such as cardiac arrhythmias are the parent to atrial fibrillation and atrial flutter. Additional phenotypes that are not represented directly in the ICD9 hierarchy are also included, such as inflammatory bowel disease as the parent for Crohn’s disease and ulcerative colitis. Diagnoses that were not possible for an individual were set to missing, such as pregnancy for males, or prostate diseases for females. Detailed features of all phenotype algorithms used are available from: http://phewascatalog.org.

### Clinical Outcomes

For each phenotype, we regressed the binary outcome onto the standardized heterozygosity from equation 2 above, adjusted for birth year and the top 5 principal components of ancestry using logistic regression. We limited analysis to outcomes with 40 or more cases and individuals with at least 2 ICD9 codes. We determined the threshold for statistical significance by Bonferroni correction for the number of analyses where the model converged.

## Data and Software Availability

The data that support the findings of this study are available from the BioVU resource, but restrictions apply to the availability of these data, which were used under license for the current study, and so are not publicly available. Data are however available upon reasonable request with permission from BioVU.

## Acknowledgements

The authors would like to thank J. Henshaw and W. Kunce for useful feedback on the manuscript. The dataset used in the analyses described were obtained from Vanderbilt University Medical Centers BioVU which is supported by institutional funding and by the Vanderbilt CTSA grant UL1 TR000445 from NCATS/NIH. This work was supported by the National Institutes of Health (grant numbers RC2GM092618, U01HG004603, K23HD000001, R25/T32CA160056, R01LM010685).

