## Supplemental Figures and Tables for "Temporal changes in genetic admixture are linked to heterozygosity and health diagnoses in humans"

**SUPPLEMENTARY INFORMATION**

**Figure S1.** Histogram showing distribution of subjects by EHR designated ancestry and birth year. The numbers of individuals by birth years studied depicting the relative fraction of individuals identified as non-Hispanic White, non-Hispanic Black, Hispanic/Latino, non- Hispanic Asian in the EHR. Samples are in one year bins in the histogram.

**Figure S2.** Principal components plots of study individuals by decades of birth between 1905-2013. Principal components were calculated in EIGENSTRAT with populations in 1000 Genomes to serve as references points for the plotting ancestry groups. PC1 is on the y-axis and PC2 is on the x-axis. Blue = non-Hispanic White, Green = non-Hispanic Black, Red = Hispanic/Latino, Gold = non-Hispanic Asian.

**Figure S3.** Proportions of ancestry derived by STRUCTURE analysis of AIMs for study individuals plotted against birth year. The y-axis has ancestry proportion as determined from STRUCTURE software while birth year is on the x-axis for the entire study population. The ancestry lines are color coded and 95% confidence intervals are indicated.

**Figure S4.** Proportions of ancestry derived by STRUCTURE analysis of AIMs for EHR-designated non-Hispanic White study individuals plotted against birth year. The y-axis has ancestry proportion as determined from STRUCTURE software while birth year is on the x-axis for the individuals designated as non-Hispanic White in the EHR.

**Figure S5.** Proportions of ancestry derived by STRUCTURE analysis of AIMs for EHR-designated non-Hispanic Black study individuals plotted against birth year. The y-axis has ancestry proportion as determined from STRUCTURE software while birth year is on the x-axis for the individuals designated as non-Hispanic Black in the EHR.

**Figure S6.** Proportions of ancestry derived by STRUCTURE analysis of AIMs for EHR-designated Hispanic/Latino study individuals plotted against birth year. The y-axis has ancestry proportion as determined from STRUCTURE software while birth year is on the x-axis for the individuals designated as Hispanic/Latino in the EHR.

**Figure S7.** Proportions of ancestry derived by STRUCTURE analysis of AIMs for EHR-designated non-Hispanic Asian study individuals plotted against birth year. The y-axis has ancestry proportion as determined from STRUCTURE software while birth year is on the x-axis for the individuals designated as non-Hispanic Asian in the EHR.

**Figure S8**. Standardized heterozygosity plotted against birth year. The y-axis depicts the standardized heterozygosity of individuals with birth year on the x-axis. Cohorts are grouped by stated race and 95% confidence intervals are indicated.

**Figure S9.** Plot of individual-level 2013 US Census data for Tennessee. The average number of race categories claimed by each individual on the y-axis against birth year on the x- axis.

**Figure S10.** Plot of individual-level 2013 US Census data for the East South Central Division. The average number of r categories claimed by each individual on the y-axis against birth year on the x-axis.

**Figure S11.** Plot of individual-level 2013 US Census data for the entire United States. The average number of race categories claimed by each individual on the y-axis against birth year on the x-axis.

**Figure S12.** Phenome-Wide Association Study results for analysis of standardized heterozygosity and clinical outcomes. A series of association tests were conducted using clinical outcomes classified from the EHRs of study subjects, adjusted for 5 PCs, sex, and birth year.

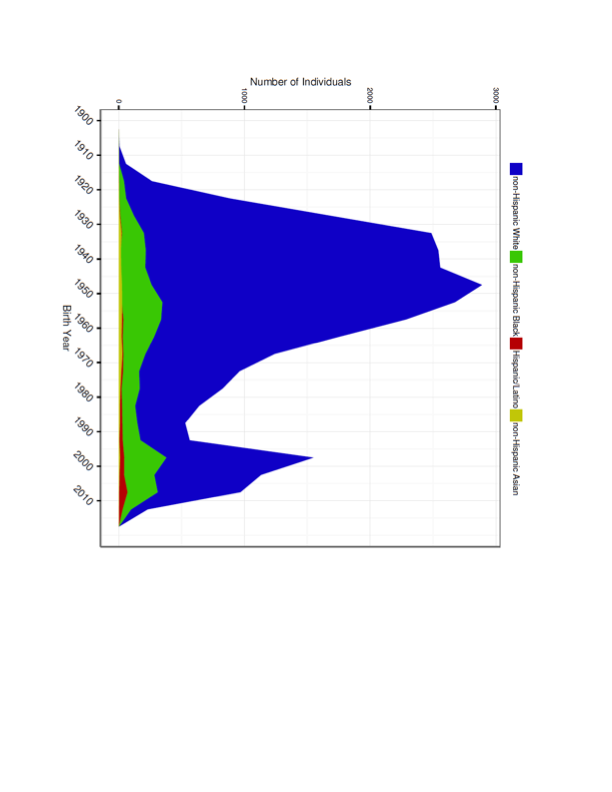

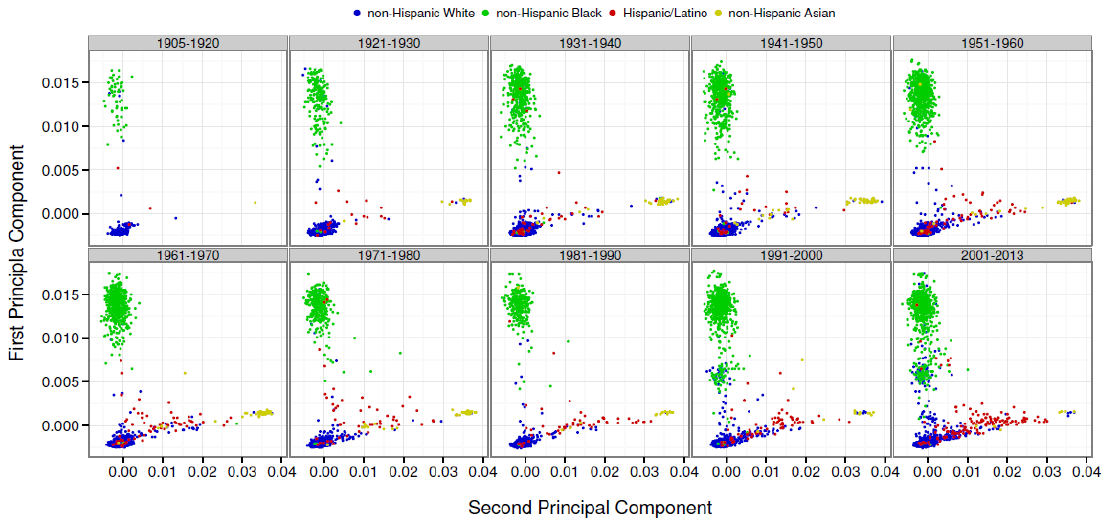

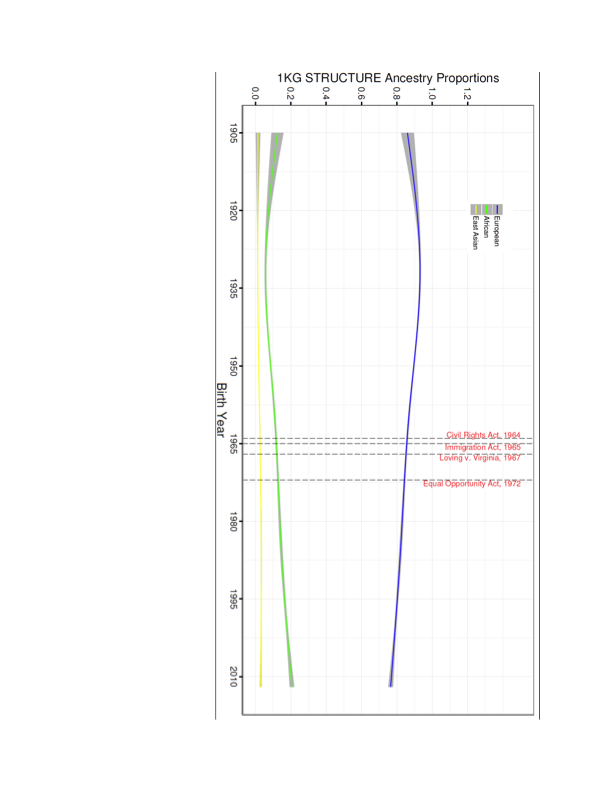

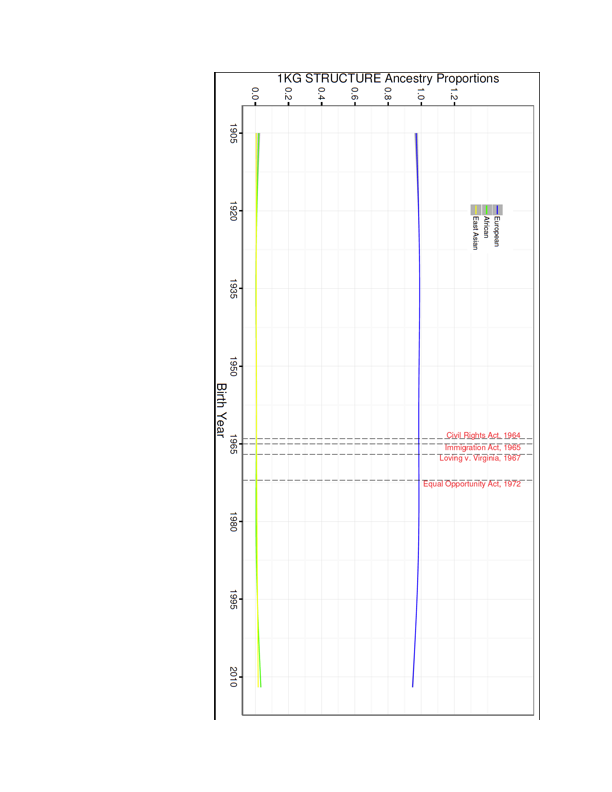

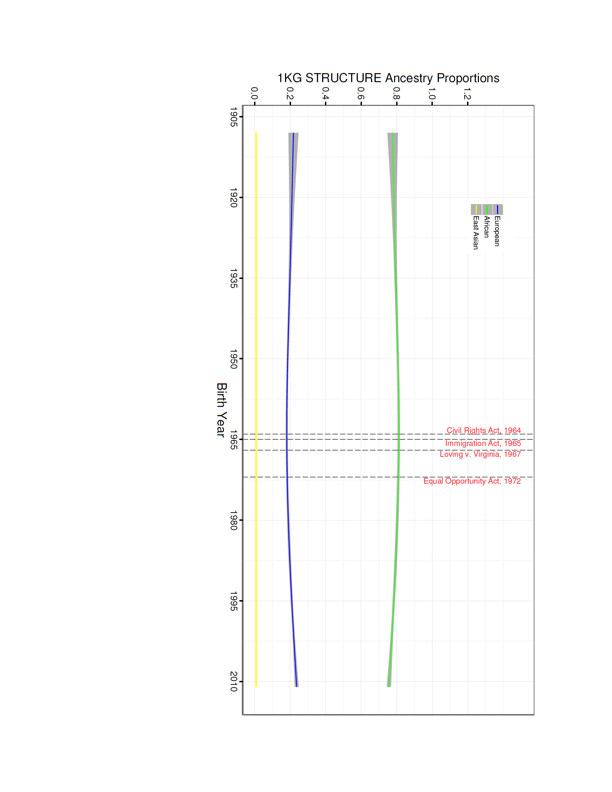

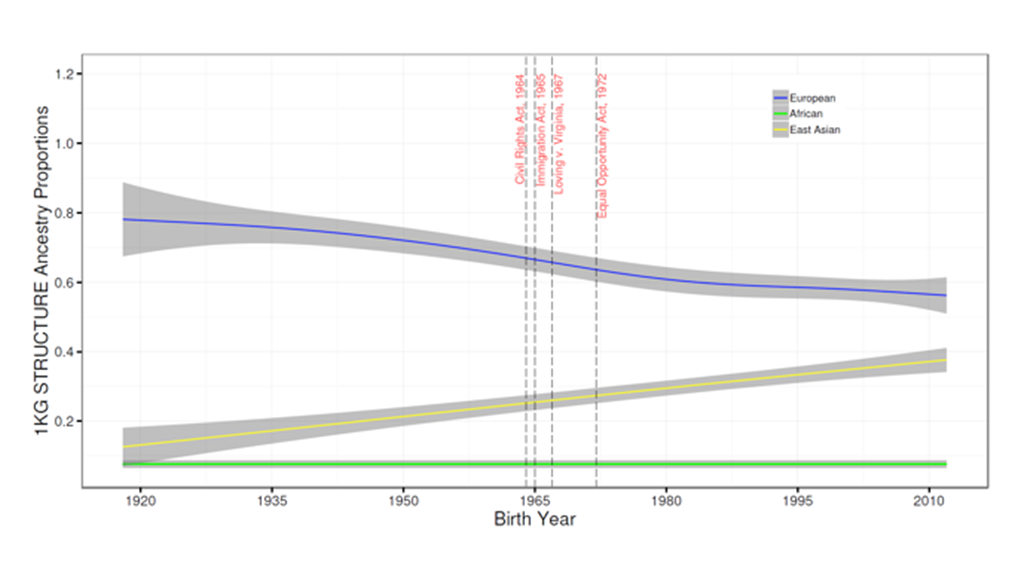

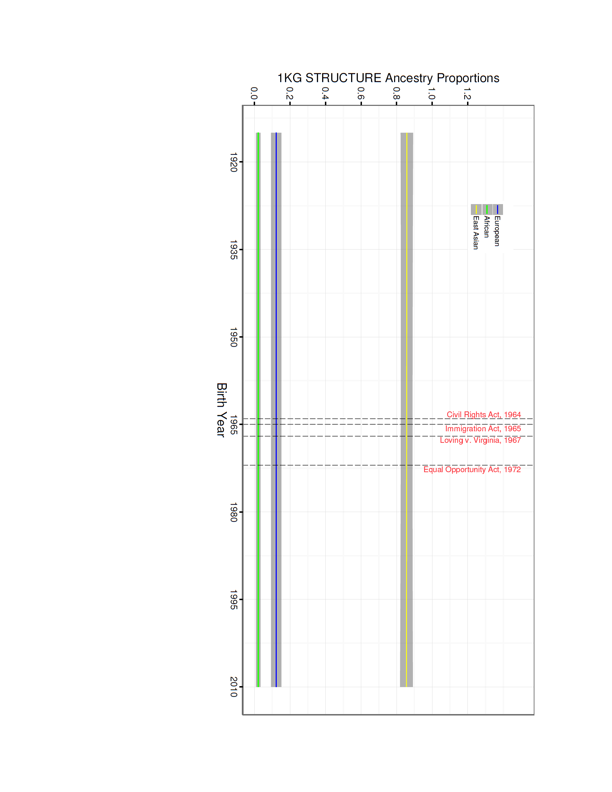

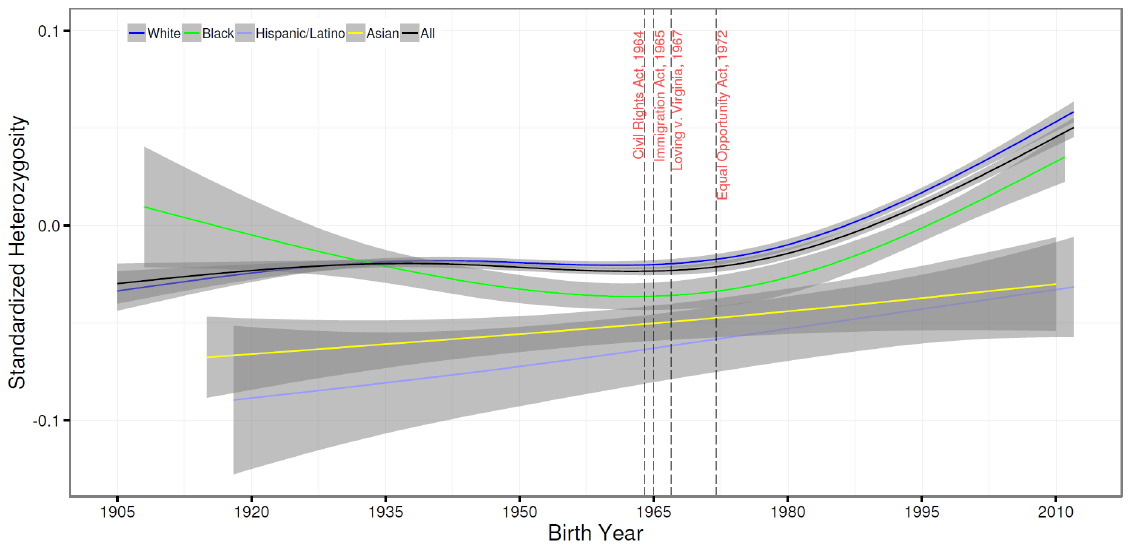

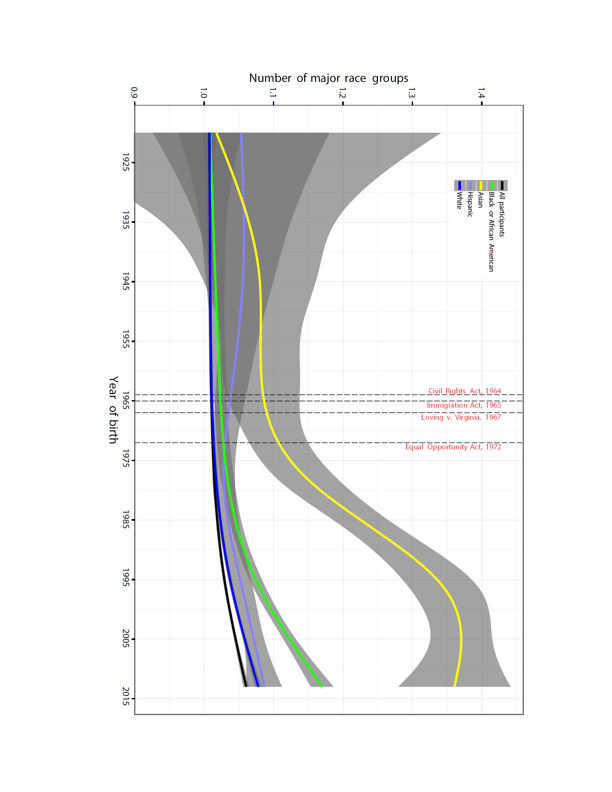

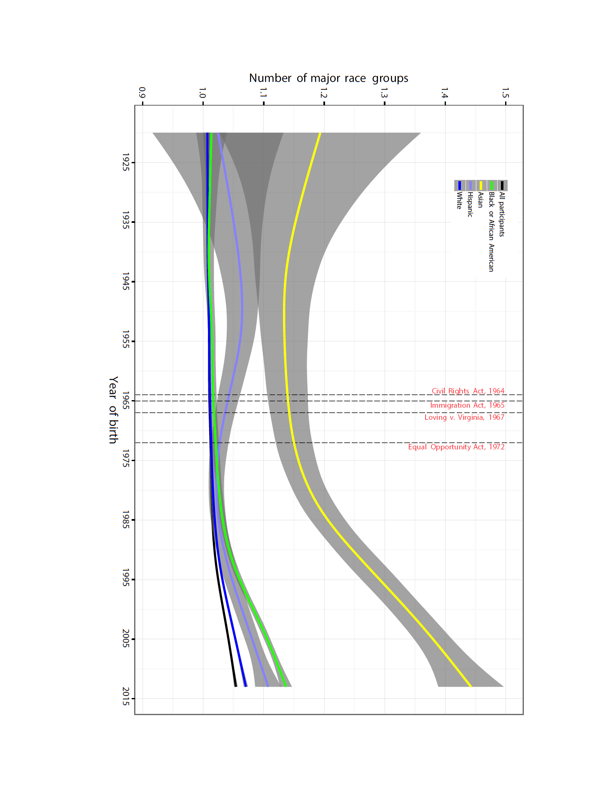

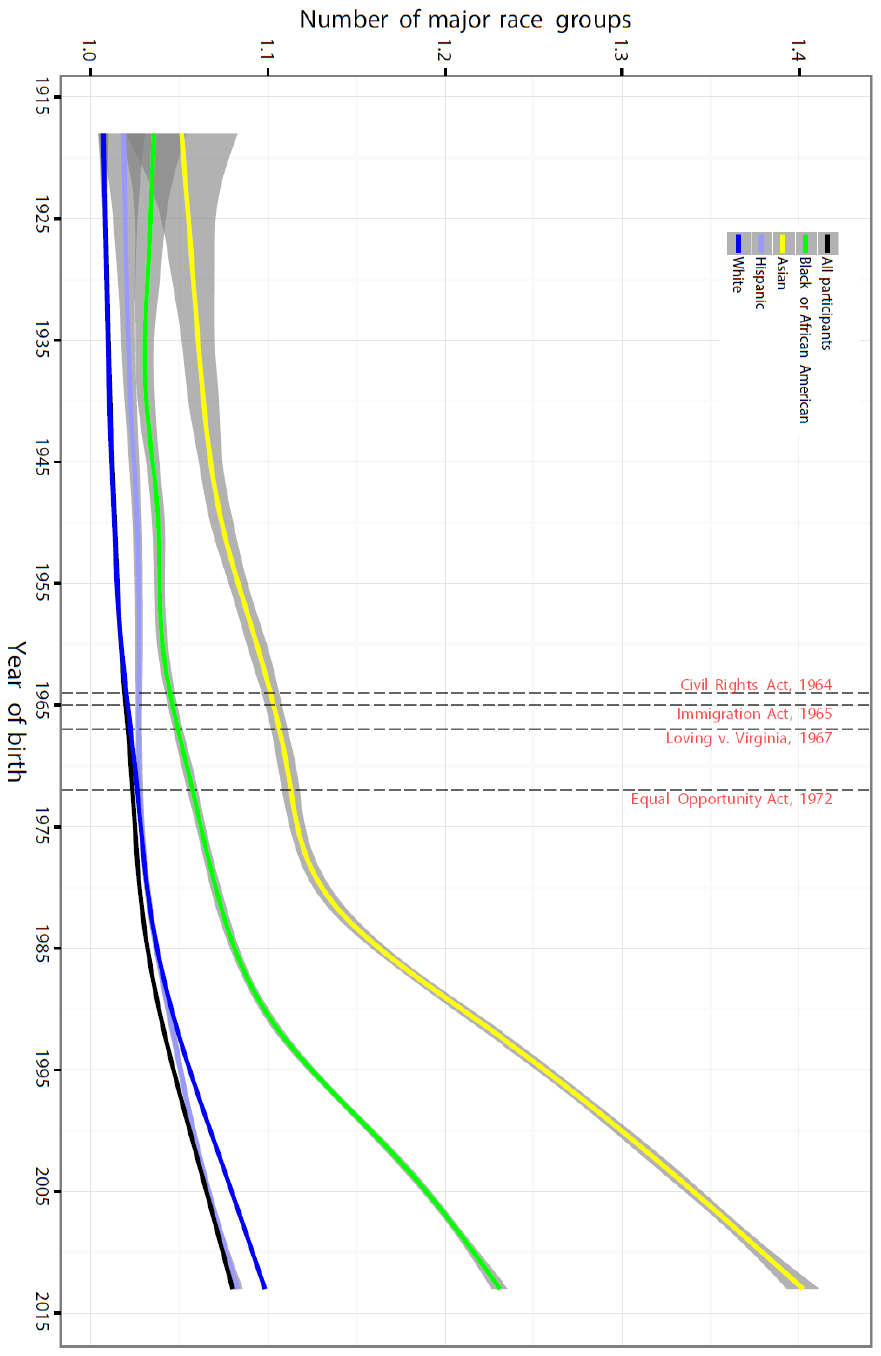

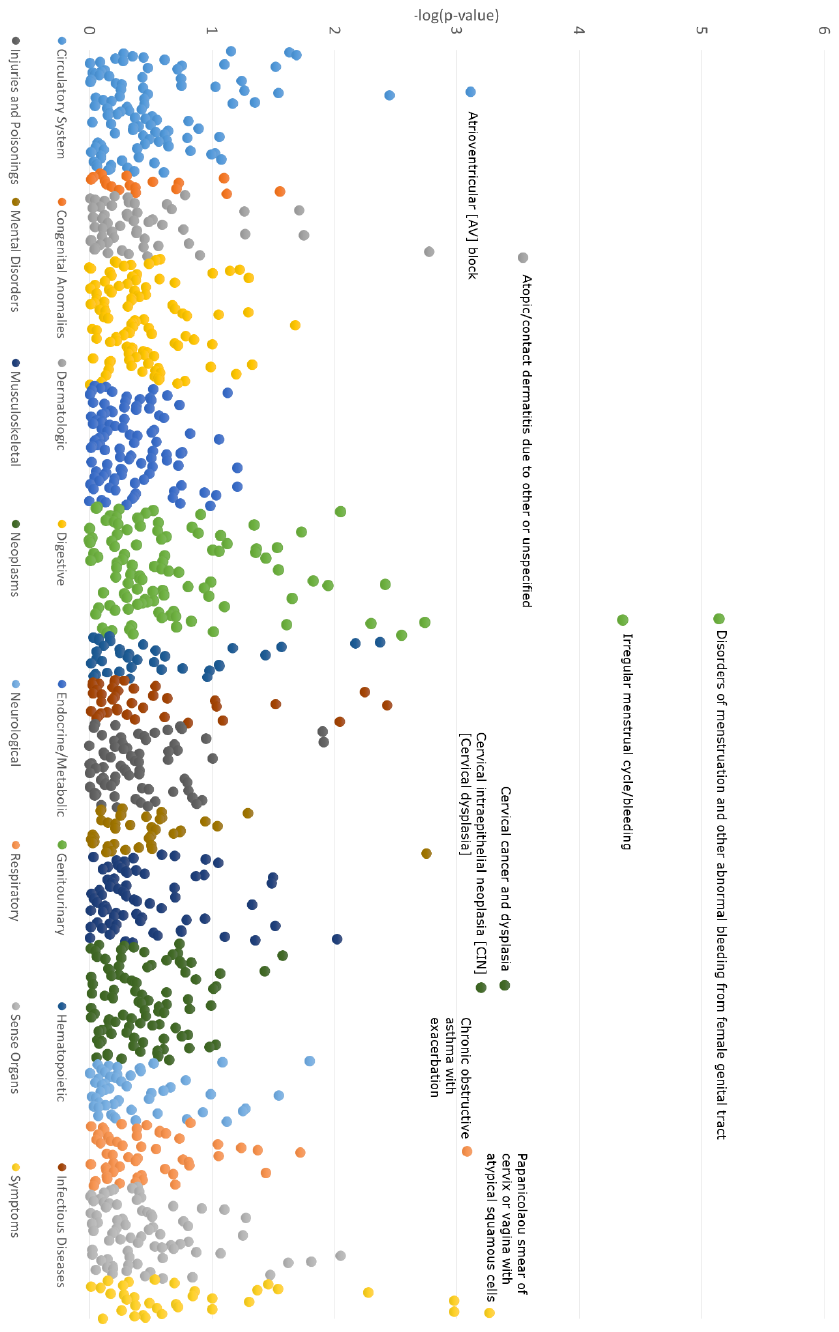

**Table S1**. Population demographics by EHR race

| **Race** | **White** | **Black** | **Hispanic/Latino** | **Asian** | **Other/Unknown^1^** |
| --- | --- | --- | --- | --- | --- |
| **N (%)** | 28,723 (80.1) | 4,129 (11.5) | 550 (1.5) | 270 (0.75) | 2,170 (2.8) |
| **Male %** | 46.9% | 38.9% | 43.6% | 42.5% | 39.6% |
| **Birth year** |  |  |  |  |  |
| **Mean (SD)** | 1957 (24.1) | 1968 (26.0) | 1976 (25.6) | 1959 (19.8) | 1955 (20.2) |
| **Median (IQR)** | 1951 (1938–1971) | 1964 (1948–1995) | 1977 (1957–2000) | 1958 (1944-1972) | 1953 (1940–1967) |
| **Range** | 1905–2012 | 1908–2011 | 1918–2012 | 1915-2010 | 1906–2012 |

^1^Other/Unknown-this category included individuals whose race was not documented in the EHR and listed as “Unknown” or “Declined” and American Indian or Alaska Native individuals. We did not include American Indian or Alaska Native due to the limited sample size (N=30).

**Table S2.** Mean admixture proportion $\pm$ SE at 15-year intervals. Admixture Proportion estimated from GAM in each race group (in Figure 1).

|  |  | Birth Year | | | | | | |
| --- | --- | --- | --- | --- | --- | --- | --- | --- |
|  | N | 1920 | 1935 | 1950 | 1965 | 1980 | 1995 | 2010 |
| White | 28723 | 0.011 $\pm$0.001 | 0.007$\pm$0.001 | 0.009$\pm$0.0004 | 0.011$\pm$0.001 | 0.011$\pm$0.001 | 0.021$\pm$0.001 | 0.042$\pm$0.001 |
| Black | 4129 | 0.207$\pm$ 0.006 | 0.196$\pm$0.003 | 0.186$\pm$0.002 | 0.182$\pm$0.002 | 0.186$\pm$0.003 | 0.199$\pm$0.002 | 0.217$\pm$0.004 |
| Hispanic/Latino | 550 | 0.199 $\pm$ 0.041 | 0.181$\pm$0.017 | 0.210$\pm$0.015 | 0.275$\pm$0.014 | 0.310$\pm$0.013 | 0.329$\pm$0.013 | 0.356$\pm$0.018 |
| East Asian | 270 | 0.048 $\pm$ 0.020 | 0.059$\pm$0.014 | 0.070$\pm$0.010 | 0.082$\pm$0.010 | 0.096$\pm$0.013 | 0.112$\pm$0.018 | 0.128$\pm$0.027 |
| All | 33672 | 0.030 $\pm$0.002 | 0.021$\pm$0.001 | 0.030$\pm$0.001 | 0.040$\pm$0.001 | 0.047$\pm$0.001 | 0.067$\pm$0.001 | 0.010$\pm$0.002 |

**Table S3**: Mean $\pm$ SE of standardized heterozygosity at 15-year birth intervals (in figure 2).

|  |  | Birth Year | | | | | | |
| --- | --- | --- | --- | --- | --- | --- | --- | --- |
|  | N | 1920 | 1935 | 1950 | 1965 | 1980 | 1995 | 2010 |
| White | 28723 | -0.024$\pm$0.002 | -0.019$\pm$0.001 | -0.019$\pm$0.001 | -0.020$\pm$0.001 | -0.010$\pm$0.001 | 0.017$\pm$0.001 | 0.053$\pm$0.002 |
| Black | 4129 | -0.005$\pm$0.009 | -0.020$\pm$0.004 | -0.032$\pm$0.004 | -0.036$\pm$0.004 | -0.027$\pm$0.004 | -0.001$\pm$0.004 | 0.033$\pm$0.006 |
| Hispanic/Latino | 550 | -0.089$\pm$0.018 | -0.081$\pm$0.013 | -0.072$\pm$0.010 | -0.063$\pm$0.009 | -0.053$\pm$0.008 | -0.043$\pm$0.009 | -0.033$\pm$0.012 |
| East Asian | 270 | -0.066$\pm$0.009 | -0.061$\pm$0.006 | -0.056$\pm$0.005 | -0.050$\pm$0.005 | -0.044$\pm$0.006 | -0.037$\pm$0.008 | -0.030$\pm$0.012 |
| All | 33672 | -0.023$\pm$0.002 | -0.0196$\pm$0.001 | -0.021$\pm$0.001 | -0.023$\pm$0.001 | -0.014$\pm$0.001 | 0.011$\pm$0.001 | 0.045$\pm$0.002 |

**Table S4**: Table S3: Mean $\pm$ SE Physical Distance at 9.5Mb associated with Figure 3

| Birth Year | Mean & SE |
| --- | --- |
| 1905-1929 | 0.094$\pm$0.0003 |
| 1930-1939 | 0.0883$\pm$0.0003 |
| 1940-1949 | 0.0887$\pm$0.0003 |
| 1950-1959 | 0.1068$\pm$0.0005 |
| 1960-1969 | 0.1167$\pm$0.0003 |
| 1970-1979 | 0.1246$\pm$0.0005 |
| 1980-1989 | 0.1402$\pm$0.0006 |
| 1990-1999 | 0.1314$\pm$0.0009 |
| 2000-2013 | 0.1332$\pm$0.0004 |

**Table S5.** Secondary analysis of top PheWAS signals in combined adult-only and adult female-only

subsets.

| **PheCode** | **Phenotype** | **Full Sample** | | **Adults only** | | **Adult Females only** | |
| --- | --- | --- | --- | --- | --- | --- | --- |
|  |  | **P-value** | **OR (95% Confidence Interval)** | **P-value** | **OR (95% Confidence Interval)** | **P-value** | **OR (95% Confidence Interval)** |
| 626 | Disorders of menstruation and other abnormal bleeding from female genital tract | 7.21x10-6 | 0.37 (0.24 – 0.57) | 3.00x10-3 | 0.48 (0.29 – 0.78) | 3.25x10-4 | 0.39 (0.23 – 0.65) |
| 626.1 | Irregular menstrual cycle/bleeding | 4.37x10-5 | 0.37 (0.23 – 0.60) | 7.24x10-3 | 0.49 (0.29 – 0.82) | 9.85x10-4 | 0.40 (0.23 – 0.69) |
| 939 | Atopic/contact dermatitis due to other or unspecified | 2.86x10-4 | 1.82 (1.32 – 2.52) | 0.048 | 1.63 (1.00 – 2.66) | 0.36 | 1.33 (0.72 – 2.46) |
| 180 | Cervical cancer and dysplasia | 4.02x10-4 | 0.19 (0.08 – 0.48) | 2.05x10-3 | 0.22 (0.08 – 0.57) | 1.57x10-3 | 0.20 (0.08 – 0.45) |
| 792.1 | Papanicolaou smear of cervix or vagina with atypical squamous cells | 5.36x10-4 | 0.21 (0.09 – 0.51) | 4.09x10-3 | 0.25 (0.10 – 0.64) | 2.33x10-3 | 0.22 (0.08 – 0.58) |
| 180.3 | Cervical intraepithelial neoplasia [CIN] [Cervical dysplasia] | 6.24x10-4 | 0.15 (0.05 – 0.45) | 2.61x10-3 | 0.17 (0.06 – 0.54) | 1.54x10-3 | 0.15 (0.05 – 0.49) |
| 426.2 | Atrioventricular [AV] block | 7.62x10-4 | 2.97 (1.58 – 5.61) | 0.81 | 1.27 (0.18 – 8.76) | 0.73 | 0.56 (0.02 – 16.43) |
| 495.11 | Chronic obstructive asthma with exacerbation | 8.13x10-4 | 4.63 (1.89 – 11.34) | 2.46x10-3 | 5.77 (1.84 – 18.09) | Failed to converge | |
| 785 | Abdominal pain | 1.04x10-3 | 0.70 (0.57 – 0.87) | Failed to converge | | 0.7 | 0.93 (0.66 – 1.32) |
| 792 | Abnormal Papanicolaou smear of cervix and cervical Human Papillo (HPV) | 1.04x10-3 | 0.33 (0.17 – 0.64) | 0.015 | 0.41 (0.20 – 0.84) | 0.011 | 0.39 (0.19 – 0.80) |
| 709.2 | Sicca syndrome | 1.66x10-3 | 4.38 (1.75 – 10.99) | Failed to converge | | 9.62x10-5 | 7.21 (2.67 – 19.45) |
| 318 | Tobacco use disorder | 1.75x10-3 | 0.48 (0.30 – 0.76) | 0.036 | 0.59 (0.36 – 0.97) | 0.58 | 0.84 (0.45 – 1.57) |
| 626.12 | Excessive or frequent menstruation | 1.80x10-3 | 0.28 (0.12 – 0.62) | Failed to converge | | 4.37x10-3 | 0.27 (0.11 – 0.66) |
| 628 | Ovarian cyst | 2.83x10-3 | 0.23 (0.09 – 0.60) | 0.02 | 0.29 (0.11 – 0.82) | 9.21x10-3 | 0.25 (0.09 – 0.71) |
| 426.24 | Atrioventricular block, complete | 3.57x10-3 | 3.30 (1.48 – 7.38) | Failed to converge | | Failed to converge | |

**Table S6**. Average pairwise Δp from 1000 Genomes Project reference populations for AIMs used in PCA,

heterozygosity, and STRUCTURE analysis.

| **AMR** | 0.51 |  |  |
| --- | --- | --- | --- |
| **ASN** | 0.46 | 0.12 |  |
| **EUR** | 0.59 | 0.09 | 0.17 |
|  | **AFR** | **AMR** | **ASN** |

AMR-Native American populations of the 1000 Genomes Project (CLM, MXL, PUR); ASN-Asian populations of the 1000 Genomes

Project (CHB, CHS, JPT); EUR-European ancestry populations in the 1000 Genomes Project (CEU, FIN, GBR, IBS, TSI); AFR-African

ancestry populations in 1000 Genomes Project (ASW, LWK, YRI)
